# FUSTr: a tool to find gene Families Under Selection in Transcriptomes

**DOI:** 10.1101/146951

**Authors:** T. Jeffrey Cole, Michael S. Brewer

## Abstract

FUSTr is a tool for finding genes in transcriptomic datasets under strong positive selection that automatically detects isoform designation patterns in transcriptome assemblies to maximize phylogenetic independence in downstream analysis. When applied to previously studied spider toxin families as well as simulated data, FUSTr successfully grouped coding sequences into proper gene families as well as correctly identified those under strong positive selection. FUSTr provides a tool capable of utilizing multi-processor high-performance computational facilities and is scalable for large transcriptomic biodiversity datasets.

**Availability:** FUSTr is freely available under a GNU license and can be downloaded at https://github.com/tijeco/FUSTr.

## Introduction

Elucidating patterns and processes involved in the adaptive evolution of genes and genomes of organisms is fundamental to understanding the vast phenotypic diversity found in nature. Recent advances in RNA-Seq technologies have played a pivotal role in expanding knowledge of molecular evolution through the generation of an abundance of protein coding sequence data across all levels of biodiversity (Todd *et al.*, 2016). In non-model eukaryotic systems, transcriptomic experiments have become the *de facto* approach for functional genomics in lieu of whole genome sequencing. This is due largely to lower costs, better targeting of coding sequences, and enhanced exploration of post-transcriptional modifications and differential gene expression (Wang *et al.*, 2009). This influx of transcriptomic data has resulted in a need for scalable tools capable of elucidating broad evolutionary patterns in large biodiversity datasets.

Billions of years of evolutionary processes gave rise to remarkably complex genomic architectures across the tree of life. Numerous speciation events along with frequent whole genome duplications have given rise to a myriad of multigene families with varying roles in the processes of adaptation (Benton, 2015). Grouping protein encoding genes into their respective families *de novo* has remained a difficult task computationally. This typically entails homology searches in large amino acid sequence similarity networks with graph partitioning algorithms to cluster coding sequences into transitive groups (Andreev and Racke, 2006). This is further complicated in eukaryotic transcriptome datasets that contain several isoforms via alternative splicing (Matlin *et al.*, 2005). Further exploration of Darwinian positive selection in these families is also nontrivial, requiring robust Maximum Likelihood and Bayesian phylogenetic approaches.

Here we present a tool for finding Families Under Selection in Transcriptomes (FUSTr), to address the difficulties of characterizing molecular evolution in largescale transcriptomic datasets. FUStr can be used to characterize selective regimes on homologous groups of phylogenetically independent coding sequences in transcriptomic datasets and has been verified using curated spider venom toxins and simulated datasets. The presented pipeline implements simplified user experience with minimized third-party dependencies, in an environment robust to breaking changes to maximize long-term reproducibility.

### 1 Implementation

FUSTr is written in Python with all data filtration, preparation steps, and command line arguments/parameters for external programs contained in the workflow engine Snakemake (Köster and Rahmann, 2012). Snakemake allows FUSTr to operate on high performance computational facilities, while also maintaining ease of reproducibility. FUSTr utilizes a custom anaconda environment that has reduced the number of third party dependencies needed by the user from seven to a mere two software packages. FUSTr takes as input assembled transcriptomes from any number of taxa in FASTA format. Header patterns are analyzed to auto-detect whether the given assembly includes isoforms by comparing the header patterns to common assemblers (Haas *et al.*, 2013) and detecting naming convention redundancies commonly used in isoform designations. Single best Open Reading Frames from transcripts are then extracted using Transdecoder v3.0.1 (Haas *et al.*, 2013), providing nucleotide coding sequences (CDS) and complimentary amino acid sequences. This facilitates further analyses requiring codon level sequences while using the more informative amino acid sequences for homology inferences and multiple sequence alignments. If the data contain several isoforms of the same gene, at this point only the longest isoform is kept for further analysis to ensure phylogenetic independence.

Homology is inferred via all against all BLASTP v2.5.0. This homology network is then grouped into putative gene families using transitive clustering with SiLiX v.1.2.11 (Miele *et al.*, 2011). Multiple amino acid sequence alignments of each family are then generated using the appropriate algorithm automatically detected using MAFFT v7.221 (Katoh and Standley, 2013). Spurious columns in alignments are removed with Trimal v1.4.1’s *gappyout* algorithm (Capella-Gutiérrez and Silla-Martínez, 2009). An approximately maximum likelihood phylogeny of each family is inferred using FastTree v2.1.9 (Price *et al.*, 2010). Multiple sequence codon alignments are then generated by reverse translation of the amino acid alignment using the CDS sequences. Tests of pervasive positive selection at site specific amino acid level are implemented on families containing at least 15 sequences using codeml v4.9 (Yang, 2007) with the codon alignments and inferred phylogeny. Log-likelihood values of codon substitution models that allow positive selection are then compared to respective nested models not allowing positive selection (M0/M3, M1a/M2a, M7/M8, M8a/M8), Bayes Empirical Bayes (BEB)

analysis then determines posterior probabilities that the ratio of nonsynonymous to synonymous substitutions (dN/dS) exceeds one for individual amino acid sites. The final output is a summary file describing what gene families were detected, and those that are under strong selection.

We tested FUSTr with seven highly curated spider venom toxin protein families comprising 624 coding sequences that were analyzed for selective regimes in previous studies (Sunagar and Moran, 2015). Detailed results and included families can be found in Fig. 1 and Supplemental File 1. FUSTr recovered all 7 families correctly. A single family with >15 members, Kunitz toxins, was found to be under strong positive selection, corroborating previous results (Sunagar and Moran, 2015).

**Figure 1.**
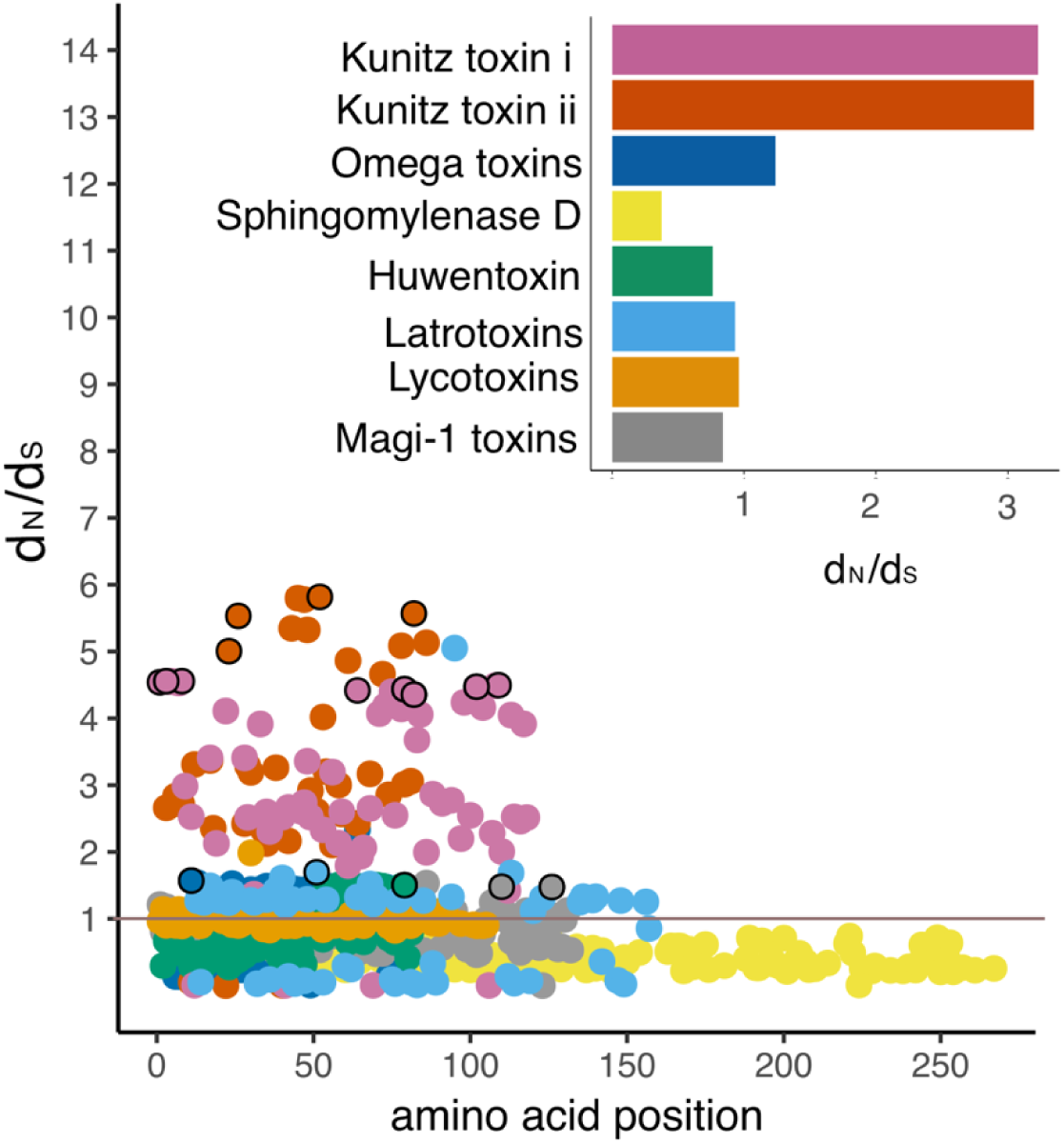
Spider toxin family positive selection. Amino acid site specific dN/dS values for families from bar plot color code. Points outlined in black represent sites under positive selection with posterior probability > 0.95. Grey line indicates neutrality threshold.

We also utilized coding sequences from simulated gene families with predetermined selective regimes to further validate FUSTr. We used EvolveAGene (Hall, 2007) on 50 random coding sequences of a random length of 50-100 codons to generate gene families containing 16 sequences each with average branch lengths chosen randomly from 0.01- 0.40 evolutionary units. All families were simulated to undergo pervasive positive selection in random amino acid sites. FUSTr successfully recovered all 50 families and identified the correct amino acid sites under positive selection in each family. Scripts for these simulations can be found at https://github.com/tijeco/FUSTr.

### 2 Conclusions

Current advances in RNA-seq technologies have allowed for a rapid proliferation of transcriptomic datasets in numerous non-model study systems. FUSTr provides a useful tool for novice bioinformaticians to detect gene families in transcriptomes under strong selection. Results from this tool can provide information about candidate genes involved in the processes of adaptation, in addition to contributing to functional genome annotation.

## Acknowledgements

This work would not have been possible without XSEDE computational allocations (BIO160060). We also thank Chris Cohen for editing this manuscript.

